# MassGenie: a transformer-based deep learning method for identifying small molecules from their mass spectra

**DOI:** 10.1101/2021.06.25.449969

**Authors:** Aditya Divyakant Shrivastava, Neil Swainston, Soumitra Samanta, Ivayla Roberts, Marina Wright Muelas, Douglas B. Kell

## Abstract

The ‘inverse problem’ of mass spectrometric molecular identification (‘given a mass spectrum, calculate the molecule whence it came’) is largely unsolved, and is especially acute in metabolomics where many small molecules remain unidentified. This is largely because the number of experimentally available electrospray mass spectra of small molecules is quite limited. However, the forward problem (‘calculate a small molecule’s likely fragmentation and hence at least some of its mass spectrum from its structure alone’) is much more tractable, because the strengths of different chemical bonds are roughly known. This kind of molecular identification problem may be cast as a language translation problem in which the source language is a list of high-resolution mass spectral peaks and the ‘translation’ a representation (for instance in SMILES) of the molecule. It is thus suitable for attack using the deep neural networks known as transformers. We here present MassGenie, a method that uses a transformer-based deep neural network, trained on ~6 million chemical structures with augmented SMILES encoding and their paired molecular fragments as generated *in silico*, explicitly including the protonated molecular ion. This architecture (containing some 400 million elements) is used to predict the structure of a molecule from the various fragments that may be expected to be observed when some of its bonds are broken. Despite being given essentially no detailed nor explicit rules about molecular fragmentation methods, isotope patterns, rearrangements, neutral losses, and the like, MassGenie learns the effective properties of the mass spectral fragment and valency space, and can generate candidate molecular structures that are very close or identical to those of the ‘true’ molecules. We also use VAE-Sim, a previously published variational autoencoder, to generate candidate molecules that are ‘similar’ to the top hit. In addition to using the ‘top hits’ directly, we can produce a rank order of these by ‘round-tripping’ candidate molecules and comparing them with the true molecules, where known. As a proof of principle, we confine ourselves to positive electrospray mass spectra from molecules with a molecular mass of 500Da or lower. The transformer method, applied here for the first time to mass spectral interpretation, works extremely effectively both for mass spectra generated *in silico* and on experimentally obtained mass spectra from pure compounds. The ability to create and to ‘learn’ millions of fragmentation patterns *in silico*, and therefrom generate candidate structures (that do not have to be in existing libraries) <u>directly</u>, thus opens up entirely the field of *de novo* small molecule structure prediction from experimental mass spectra.

## Introduction

The measurement of small molecules within biological matrices, commonly referred to as metabolomics^1,2^, is an important part of modern post-genomics. Spectrometric methods are key to their identification. For the deconvolution of matrices such as human serum, chromatography methods coupled to mass spectrometry are pre-eminent^3,4^. In mass spectrometry (MS), molecules are ionised and enter a gas phase, commonly using electrospray methods, and the masses and intensities of the fragments – the mass spectrum – contains the diagnostic information that in principle represents a fingerprint for identifying the target molecule of interest. The molecular ion and its fragments may themselves be further fragmented (tandem-MS) using different energies to increase the discriminating power. Identification is currently largely performed by comparing the peaks in the mass spectra obtained with those in a library of mass spectra from known molecules, and the identities may be confirmed by running authentic chemical standards, if available.

The problem with all of the above is that the number of small molecules of potential interest (‘chemical space’) is vast^5^ (maybe 10^60 6,7^) whereas the number of synthesised and purchasable molecules as recorded at the ZINC database^8^ (http://zinc15.docking.org/), are just some 10^9^ and 6.10^6^, respectively (even most of the simplest heterocyclic scaffolds have never been made^9^). Those molecules with available, experimentally determined, high-resolution mass spectra are in the low tens or hundreds of thousands. Consequently, the likelihood that any molecule detected in a metabolomics experiment is actually in a library (or even close to a molecule that is^10^) is quite small, and most are not^11–13^. Experimentally, commonly just 10% of molecules can be identified from what are reproducible spectral features in complex matrices such as serum (e.g.^14–18^), despite the existence of many heuristics^19–22^. Solving the mass spectral molecular identification problem is thus seen widely as the key unsolved problem of metabolomics^4,11,17,23–36^. It was also seen as a classical problem in the early development of ‘artificial intelligence’^37–40^.

It is possible to compute (and hence to generate) all reasonable molecules that obey valence rules and that contain just C, H, O, N, S and halogens, which for those with C-atoms up to 17 amount to some 1.66.10^11^ molecules^41^. However, much of the problem of navigating chemical space in search of molecules that might match a given mass spectrum comes from the fact that chemical space is quasi-continuous but molecules are discrete^42^. As part of the revolution in deep learning^43,44^, *de novo* generative methods have come to the fore (e.g.^42,45–55^). These admit the *in silico* creation of vectors in a high-dimensional ‘latent’ space (‘encoding’) and their translation from and into meaningful molecular entities (‘decoding’). Although the mapping is not at all simple, small movements in this latent space from a starting point do lead to molecules that are structurally related to those at their starting points^42,56^.

By contrast to the ‘inverse problem’ that we are seeking to solve (where we have a mass spectrum and seek to find the molecule that generated it), the ‘forward problem’ of mass spectrometry is considerably more tractable (especially for high-energy electron impact mass spectra^57^). Given a known chemical structure, it is possible to fragment the weakest bonds *in silico* and thereby generate a reasonably accurate ‘mass spectrum’ of the fragments so generated (e.g.^17,58–66^). Importantly, the advent of high-mass-resolution spectra means that each mass or fragment can be contributed by only a relatively small number of (biologically) feasible molecular formulae. Even fewer can come from different, overlapping fragments. Experimentally, we shall assume that the analyst has available a mass spectrum that contains at least 10 fragmentation peaks (probably obtained using different fragmentation energies) including the protonated molecular ion, and we here confine ourselves to analysing positively charged electrospray spectra.

Part of the recent revolution in deep learning^43,44^ recognises that in order to ‘learn’ a domain it is necessary to provide the learning system with far more examples than were historically common, but that the features of (even unlabelled) images^67^ or text^68,69^ can then be learned effectively. Transformers^70,71^ are seen as currently the most successful deep learning strategy for ‘sequence-to-sequence’-type problems such as language translation^72–74^. Obviously SMILES strings constitute a language and while it may be less obvious that mass spectra do so, they too can be seen to represent a sequence of elements, in this case high-resolution peaks. The ‘spectrum-to-structure’ problem may thus be cast as a language translation problem. In particular, much as how human infants learn from their surroundings, it has been established in the transformer type of architecture^70^ that the initial ‘pre-training’ can if necessary be carried out in an entirely unsupervised manner, with ‘fine-tuning’ being sufficient to learn the domain of interest^68,71,72,75,76^. We recognise that this is also possible for the space of small molecules, where the features of interest are their (mass spectral) fragments. However, in this case we may in fact generate matched pairs of spectra and molecular structures from the encoding (as their SMILES strings) of the relevant molecules, and treat this as a supervised learning problem.

We thus considered that one might fruitfully combine these two main elements (molecular generation and molecular fragmentation) with an objective function based on the closeness of the predicted mass spectrum to the experimental one that one is trying to fit. This would allow us to ‘navigate chemical space intelligently’^52^ so as to provide the user with a set of molecules (ideally one) that generates the something close to the observed mass spectrum and hence allows identification of a candidate molecule. We here implement this approach, that we refer to as MassGenie, for learning positive electrospray ionisation mass spectra as input and the molecules that might generate them (as output). MassGenie offers an entirely novel and very promising approach and solution to the problem of small molecule identification from mass spectra in metabolomics and related fields. We believe that it represents the first use of transformers in molecular identification from (mass) spectra.

## Methods

### Small molecule datasets used

We first create and combine two datasets, that we refer to as ZINC 6M and ZINC 56M. The ZINC 6M dataset originally was a mixture of 1381 marketed drugs^77^, 158,809 natural products^78^, 1112 recon2 metabolites^77^, 150 fluorophores, and molecules from a subset of the ZINC15 database (subset1 contained 2,494,455 and subset2 (“2D-clean-drug-like-valid-instock”) contained 6,202,415 molecules. After canonicalization of the SMILES and the removal of duplicates we ended with 4,774,254 unique SMILES. We filtered any duplicates as well as structures for which the m/z of the protonated molecular ion exceeds 500. This is because comparatively few molecules had peak (mass) values higher than 500 and including them would have led to a drastic increase in the input dimensionality of our model, constraining the adjustment of other hyperparameters such as batch size, model size, etc away from their optimal values. From this, we obtained ~4.7M molecules encoded as SMILES. These will be given in published Supplementary Information. Despite these large numbers, some atoms are under-represented. For instance, we did not feel that there were sufficient S-containing molecules to give us confidence with the experimental mass spectra, although the system performed well with those generated *in silico*.

| Data source | Recon 2 | Natural products | Marketed drugs | Fluorophores | ZINC15-subset1 | ZINC15-subset2 | Total |
| --- | --- | --- | --- | --- | --- | --- | --- |
| Original smiles count | 1,112 | 158,809 | 1381 | 143 | 249,455 | 6,202,415 | 6,613,315 |
| Canonical smiles count | 1,056 | 148,971 | 1357 | 115 | 247,804 | 4,425,891 | 4,825,194<br>(4,774,258-unique) |

For ZINC 56M, we randomly sampled 56M SMILES from a set of ~794M within ZINC15 tagged as “2D-clean-druglike-annotated”. After canonicalization and duplicate removal, we obtained ~53M SMILES and again from these sampled randomly ~3.2M structures and applied the same procedure. In this purposely chosen subset of ZINC, the number of molecules with molecular ion peak values more than 500 were much lower (21,454); there were also far fewer duplicates. We then combine both datasets. The combined dataset thus had ~7.9M SMILES strings augmented to be in triplicate, of which 100,000 were reserved as a validation set (these happened to include cytosine, see later).

In addition, for fine-tuning, we made use of 201,858 molecules from the GNPS dataset (obtained from https://gnps-external.ucsd.edu/gnpslibrary/ALL_GNPS.json). The dataset originally had 439,996 data samples. We then filtered the dataset for fine tuning based on the following: first, we drop the rows containing “N/A” values. Then we filter out the samples that do not have “positive” ion mode. This left us with 271,497 samples in the dataset. After this, we applied each of the three filters that we applied in the training set, i.e. the removal of samples with m/z values exceeding 500, of samples with SMILES string length more than 99, and of the samples where the number of m/z peaks is more than 100. After this, we finally obtain a refined trainable dataset of 201,858 molecules. However, not all of these 201,858 molecules were used for training as 10,000 samples were held out as another potential test set.

We also produced a second completely independent test set of over 1000 separate molecules from ZINC 56M, fragmented as below, that were tested single-blind. Finally, we utilised experimental mass spectra generated as part of our current metabolomics studies^16,79^

### *In silico* fragmentation method

To fragment molecules *in silico*, a bespoke implementation of MetFrag^60,61^ was developed, allowing for the generation of theoretical fragmentation spectra which were tailored to approximate those found in experimental mass spectra libraries. Like MetFrag, our implementation (FragGenie, available at https://github.com/neilswainston/FragGenie) utilises the Chemistry Development Kit^80^ to represent small molecules as a matrix of atoms and bonds. Each bond is broken serially with each bond break resulting in either two fragments, or a single fragment in the case of ring-opening breaks. This process continues recursively, with each fragment being subsequently fragmented further until a specified recursion depth or a minimum fragment mass is achieved. A number of adduct groups may then be applied (e.g., protonation and sodiation) with the resulting collection of fragment m/z values providing a theoretical fragmentation spectrum for a given small molecule. (In addition, and largely for the purposes of debugging, chemical formulae and a list of bonds broken may also be generated for each individual fragment.) Such a naïve approach of breaking all bonds recursively quickly generates an excessive number of potential fragments. As such, the method was tuned against existing fragmentation spectra in the MoNA database (https://mona.fiehnlab.ucdavis.edu) in order to limit the nature of bond breaks to those which are most often found in measured spectra. While the fragmentation method of FragGenie can be filtered to include or exclude any specified bond type, for this work fragmentation was limited to single, non-aromatic bonds, protonated adducts, and a fragmentation recursion depth of 3.

Note that throughout we are in effect using only individual (not tandem) mass spectra, though these may be constructed *in silico* from experimental data by combining mass spectra generated at different fragmentation energies (e.g. by changing cone voltages in an electrospray instrument or by varying the pressure of collision gases). We also assumed that ‘real’ (experimental) mass spectra would contain a known, protonated molecular ion (reflecting a positive electrospray experiment) of a mass of sufficient accuracy to provide a molecular formula. Thus, we do not directly implement tandem spectra: ‘fragments of fragments’ are seen simply as having the molecular ion of the largest peak and may be treated accordingly in a similar manner to that described for the original molecule. This strategy necessarily misses fragments that undergo rearrangements and unknown mass losses, etc.

The fragmentation method used in our program does not seek to predict intensities directly, but merely the presence or absence of a peak with a certain high-resolution mass (m/z). We next apply a novel data augmentation procedure to make the model learn to prioritise the input peaks, to encourage the learning of peaks whose presence tended more strongly to suggest the corresponding SMILES. We observe from the MetFrag algorithm that certain types of bonds are broken in the first step. These are the ones that correspond to the highest intensity when compared with the peak list obtained from a real mass spectrometer. Also, these peaks have the same mass accuracy both theoretically as well as in the real mass spectrometer. Intuitively, this list of peaks always has (and is taken to have) the molecular ion. Subsequently, in the 2nd step, peaks with comparatively lower ranges of intensity are obtained and also with slightly higher noise in them when compared with the real peaks from the mass spectrometer. The pattern is repeated once more.

Thus, for each SMILES molecule, we generate three data samples or molecule peak pairs, one for each step of bond breaking up to three. This technically leads to a 3-fold increase in our overall data. However, in our experiments we only consider samples with a maximum input length of 100. Therefore, any SMILES molecule-FragGenie peaks pair, where the length of the list of peaks is more than 100, is dropped from the dataset. This leaves us with the ~21M SMILES that we use for training our model.

Throughout we assume that we have high-resolution mass spectral data with a precision of better than 5ppm, as is nowadays common (and is easily achieved in our own Orbitrap instruments^16^). The present paper utilises only ES+ data. For the input mass spectral data we applied a binning procedure. Data were filtered such that the peaks have an upper bound of m/z 500. This effectively produces a fixed precision of 0.01. Even for a mass of 500, a precision of 5ppm equates to a Δmass of only 0.0025, so these bins are adequate.

Thus, we consider the binning range from 0 to 50,000 and to categorize the peak into these peaks we multiply the peaks by 100 so what we get is essentially its bin number. Afterwards, the number of bins is projected as the dimensionality of the input, and the category value for every peak which we get from carrying out the above procedure reflects the index which is set in its input vector.

The input data consist of the list of values of peaks in floating point numbers. We constrained the number of peaks to be less than or equal to 100. Note that we do not use abundance data, because they depend entirely on the strength of the collisions; instead, we assume a series of experiments using collisions (or other molecular deconstruction near the source) sufficient to provide at least 10 peaks from the molecule of interest.

For the output data, we first split the SMILES into a list of individual atom types (C, N, O, P, etc) and other characters such as “[“, “]”, “(“,”)”, “=”, “-”, etc. The total vocabulary size was 69 including the 4 special tokens <sos> (start of sequence), <eos> (end of sequence), <unk> (unknown), and <pad> (padding) and this defines the output dimensionality of our transformer model as now described fully.

### The transformer model

We here used the standard transformer model^70^ and cast the problem as a peaks-to-SMILES translation problem (Figure 1A). Our transformer encoder, written in PyTorch, is composed of 12 transformer encoder layers plus 12 transformer decoder layers. The output dimensionality of all the sub-layers, as well as the embedding layers in both encoder and decoder, was set to 1024. Besides this, each Multi-Head attention layer of our transformer model was formed of 16 attention heads in total. We used dropout^81^ heavily; its value for the complete transformer model was set to 20%. Lastly, the dimensionality of the feed-forward layer present inside all of the transformer encoder and decoder layers, and applied to every position individually, was set at 4096. Effectively, we allow the model to see the entire input sequence containing peaks and impose a target attention mask in the decoder stack to prevent it from being exposed to the subsequent positions. Altogether, our model consisted of ~400M trainable parameters.

**Figure 1.**
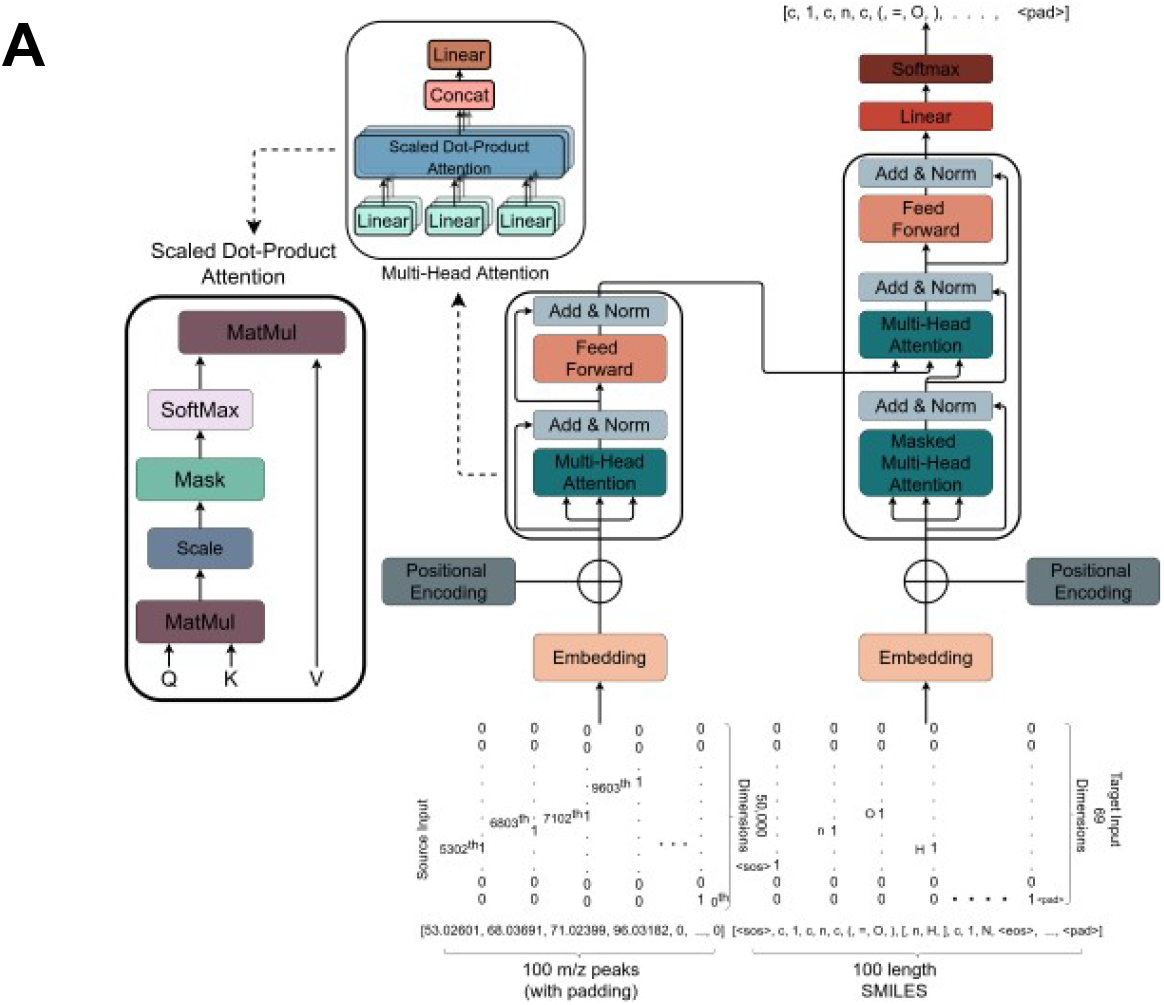
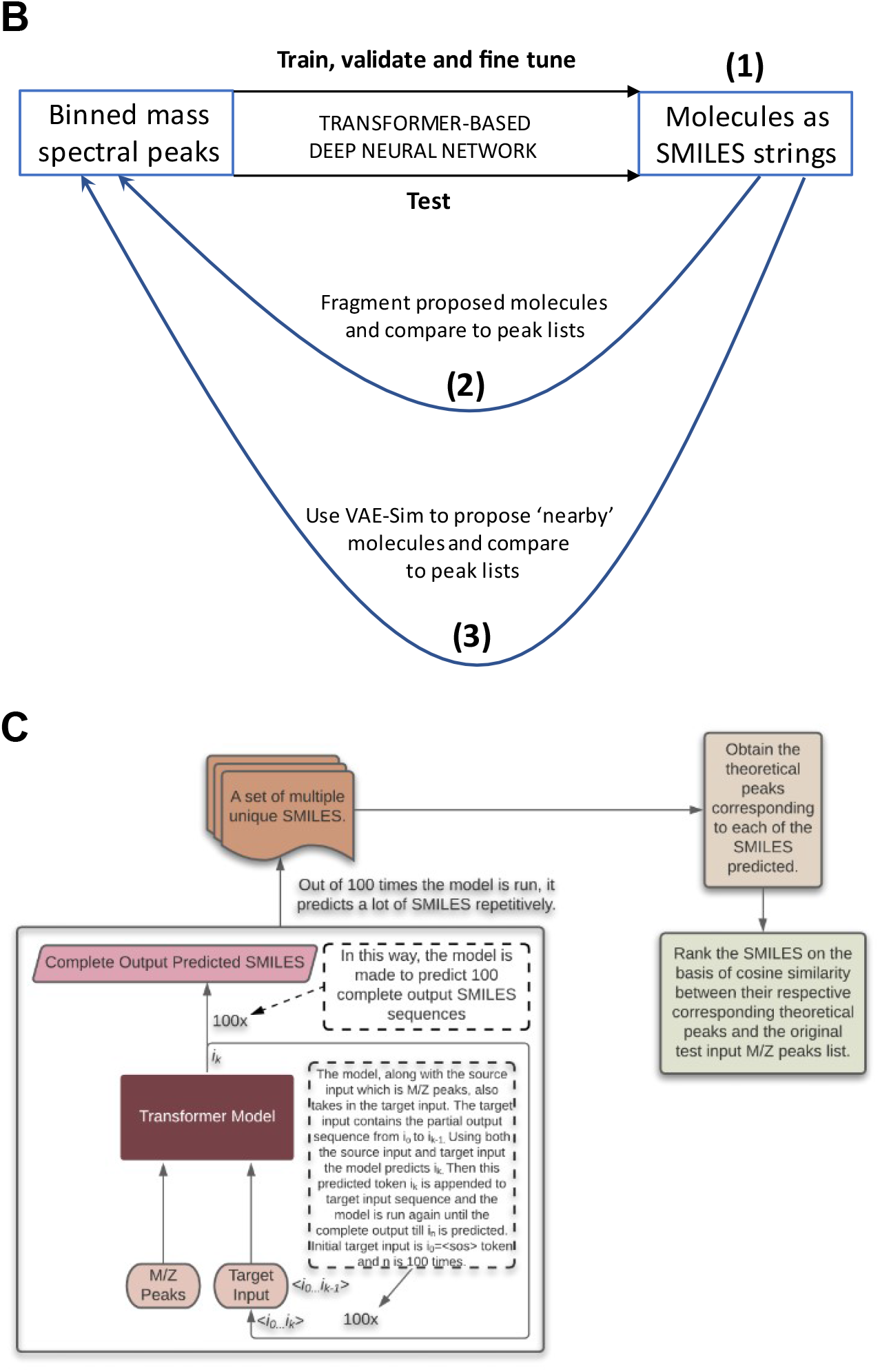
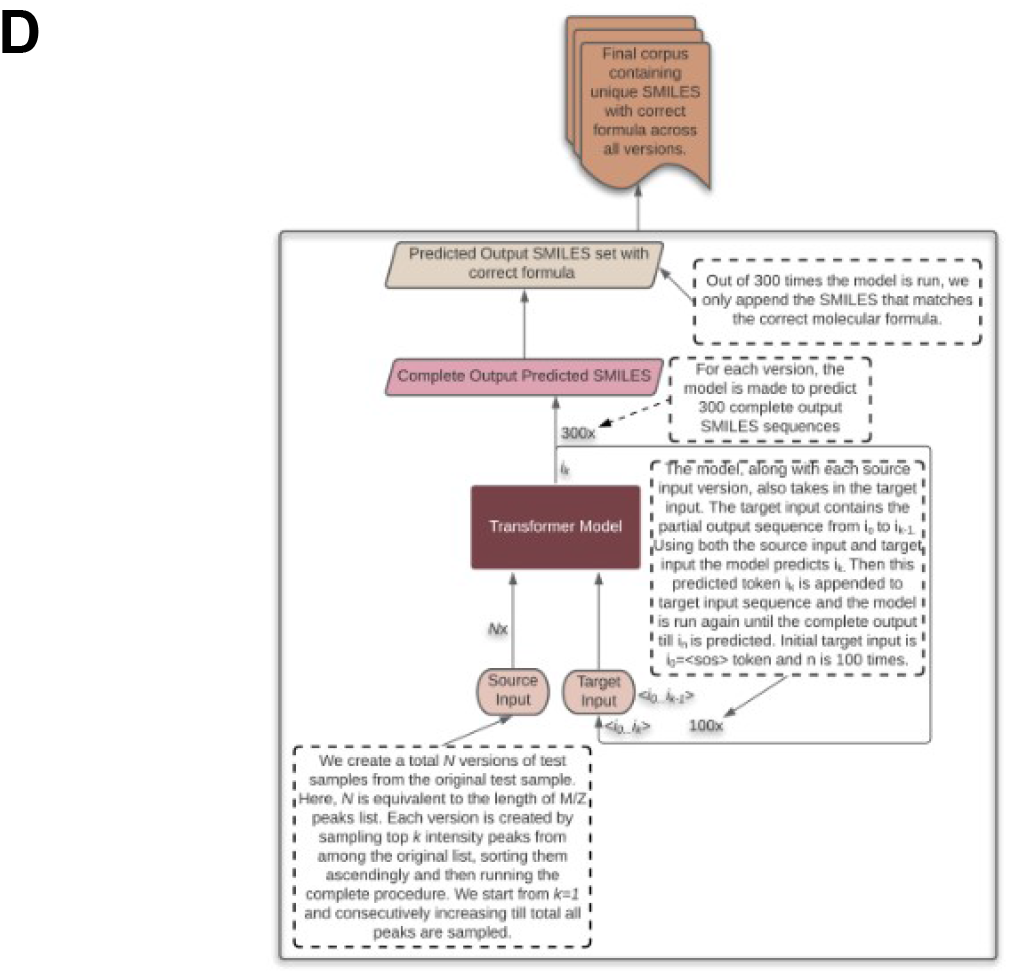
The overall strategy behind MassGenie. **A**. The overall architecture and structure of the transformer used in MassGenie, our deep learning system for identifying molecules from their mass fragments (spectra). **B**. The three basic strategies available to us for relating mass spectral peak lists to the molecules whence they might have come. (1) The transformer outputs only a single molecule. (2) We can take a series of candidate molecules, generate candidate mass spectra *in silico* using FragGenie, and compare them with the experimental mass spectra, using cosine similarity to rank order the candidates. (3) We can use VAE-Sim to generate further candidate molecules that are ‘close’ in chemical space (and possess the correct molecular formula) and rank those as in (2). This is done for both FragGenie-generated spectra (**C**) and experimental mass spectra (**D**).

As indicated, we use the transformer model, which is originally a language translation model, to ‘translate’ MS2 peak lists into SMILES. When used for language translation problems, and when given an input sentence, such a model, tends to predict multiple sentences that could be the corresponding translation of the input sentence. Some of the predicted candidate sentences even have a very similar meaning. Hence, when we apply the transformer model to our problem, we evoke a similar kind of behaviour (albeit in a different domain or context): our model predicts different SMILES strings each time when we run the model on the particular list of peaks. All of these are considered to be the candidate predictions of the molecules. Now, from our experiments and observations, what we have witnessed is when the model is pretty ‘confident’ about its predictions, it does predict the same output multiple times. However, the unique predictions are considerably smaller than the number of times it was made to predict. Also, there is typically a very high similarity between the candidate predictions. By contrast, when the model is unsure or cannot decipher the patterns properly, the model gives a set of unique predictions that is almost equal in size to the number of times the model was run for predictions. Also, the candidate predictions tend to have a considerable amount of dissimilarity between them. This admits, qualitatively, a measure of the certainty in the prediction from a transformer model. We also use the information about whether the predictions have the correct molecular formula.

### Training settings

We first trained a model with *in silico* fragments only. By applying the data augmentation described above, we used a total of ~21M data samples, from which we reserve 100,000 samples for validation and utilize the rest for training. The data were batched as 896 samples per batch. We adopt the same optimizer setup as implemented in the original paper^70^, namely the Adam optimizer with β1 = 0.9, β2 = 0.98 and ε = 10^-9^, but with the following slight changes. First, we double the warmup steps to 8000. Secondly, we apply scaling (as in^82^), to the overall learning rate, dependent upon the batch size considered for training. We use the open-source PyTorch library to construct and train the complete model for the problem. We trained our model on an NVIDIA DGX A100 system; this required approximately one day of training to reach its best validation loss. This is about 30-fold quicker than a 4-GPU (V100) machine that we have used previously^56^ for a different transformer-based deep learning problem.

### Testing the results

As with any mass spectral prediction program, the user anticipates being presented with a list of candidate predicted molecules, if possible in a rank order, and this is what we supply. It is then, of course, up to the user to validate the predictions, most persuasively^83^ by running the standard under the same conditions that generated the original mass spectrum. We employ two procedures to analyse the performance achieved by the trained model. The first procedure (“transformer search”) uses solely the transformer. When queried with a testing input list of peaks, we run the model 100 times to predict a list of output SMILES from the list, and filter this to produce a non-redundant set of predicted SMILES, also in some cases filtering to remove any molecules with the incorrect molecular formula (from the predicted molecular ion). We also have access to round tripping procedures, using our fragmentation system to regenerate the theoretical list of MS peaks corresponding to each of the non-redundant predicted SMILES. This allows us to compare the SMILES based on the similarity between the theoretically generated list of peaks and the original query list of testing peaks.

In a second procedure (“VAE search”), we use predictions from the transformer to search the local latent space in an updated version of our trained variational autoencoder^56^. We take the top-1 predicted molecule and produce a new molecule by moving a small Euclidean distance away. This is done by taking randomly 10,000 samples within a annular fixed radius from the predicted molecule’s position in the 100-dimensional latent space^56^. We also varied the radius 50 times with a fixed size of 10^-2^. This can generate many molecules beyond those on which the variational autoencoder was trained^42,56^ Out of those 500,000 samples typically ~0.1 % are valid samples (due to the effectively infinite size of the VAE latent space compared to the finite number of locally valid molecules). This takes approximately 4 min, whereupon a set of molecules with the correct mass ion is chosen.

Experimental mass spectra were generated from pooled human serum or from chemical standards using an Orbitrap mass spectrometer and the methods described in^16,79^ with standard settings, and were visualised using Compound Discoverer (Thermo).

Testing our system using ‘real’ (experimental) mass spectral peaks is somewhat different from testing it on the peaks generated *in silico*, for a number of reasons. First, we do not know exactly which peaks are more ‘important’ or even which may be true peaks, and which are erroneous due to contaminants. However, empirically, we do recognise that the higher intensity mass peaks tend more usually to be correct ones; as the intensity decreases, the noise in the peaks starts to increase and they eventually become erroneous (i.e. they do not come from the target molecule of interest). Based on this, we initiate a variable k, firstly with an initial value of 1. Then, we sample the top k intensity peaks. We sort them so that the peaks are always in an ascending order of m/z. We run the model for N=300 times and obtain a list of predictions. Since we already know the molecular formula from the exact mass of the molecular ion, the predicted list of SMILES molecules is filtered as required to match the known chemical formula. We do multiple iterations of this process, increasing the value of k by one. The number of iterations equals the total number of peaks in the test sample. Finally, we output the final list of candidate predictions. Because of the relative paucity of S- and P-containing molecules in our training set, we confined our analysis of experimental spectra to those containing solely C, H, O and N.

## Results

The first task was to train the deep transformer network described in Methods with paired *in silico* mass spectra as inputs and molecular structures as outputs. The network so generated (the code will be made available via the Supplementary Information) had some 400M nodes, and was trained over a period of ~ 1 day. We used a validation set of 100,000 molecules and use the performance (validation loss) achieved on that set as a metric to ascertain that we use the best version of the trained model for the testing. After every epoch, we run the model on the validation set and record the performance (validation loss) on it. And if the best validation loss so far is found, we record the weights of the model for that epoch. We repeat this process after every epoch until a significant number of epochs have passed without seeing any improved validation loss. The transformer architecture is given in Fig 1A, while the overall procedure including the two ‘round tripping’ strategies is given in Fig 1B. Figure 1C and 1D illustrate the strategy as applied to FragGenie-generated peaks and to experimental mass spectra.

By way of example, the following peak list was generated by FragGenie from the molecule shown on the left of Fig 2, while the candidate predictions are given on the right-hand side of that Figure.

[60.02059, 66.02125, 70.00248, 81.03214, 81.04474, 96.05563, 109.99178, 112.04313, 116.018036, 125.002686, 127.05403, 127.91178, 131.02895, 131.04152, 134.03624, 135.01643, 137.9867, 146.05243, 149.04715, 150.02734, 150.03992, 152.9976, 153.98161, 165.05083, 168.99251, 201.92743, 205.04015, 216.93832, 229.0846, 229.92235, 233.03505, 236.89629, 239.06648, 244.93324, 245.91727, 249.02995, 251.90718, 255.03694, 259.01187, 260.9282, 264.8912, 267.0614, 274.03534, 279.08142, 279.9021, 283.03186, 283.05634, 287.00677, 295.05185, 296.9758, 299.02676, 302.03024, 303.00168, 316.92114, 324.97076, 331.94464, 340.96567, 340.9657, 344.9161, 346.97266, 350.94757, 359.9396, 360.91104, 362.94308, 365.97107, 366.918, 374.9676, 378.9425, 381.9415, 385.9164, 386.98758, 390.93802, 390.9625, 390.96252, 393.966, 394.91293, 394.93744, 400.9399, 402.958, 406.93292, 406.93295, 409.93643, 410.90787, 413.91135, 428.93484, 444.92978, 459.95325]

**Figure 2.**
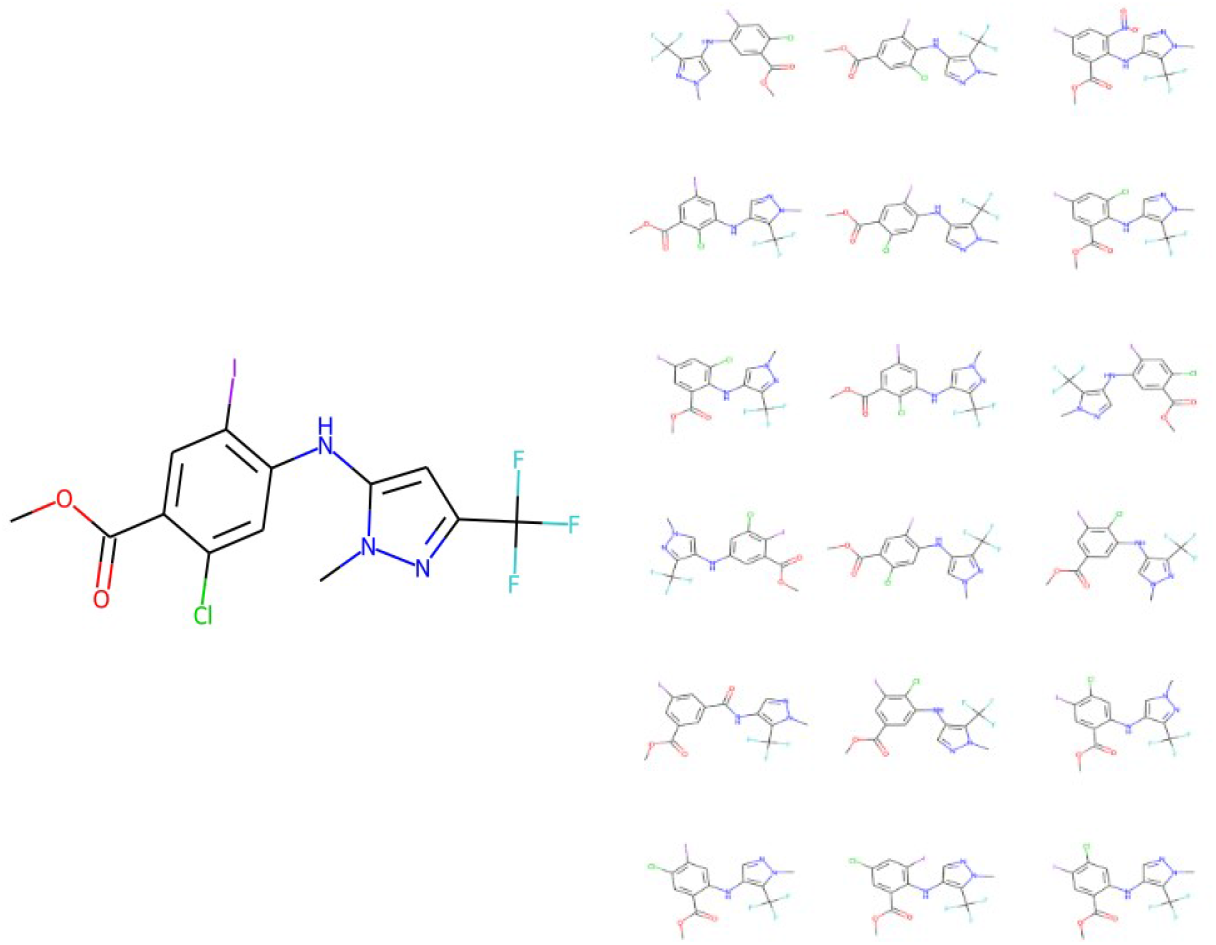
Illustration of the means by which MassGenie can predict a series of candidate molecules. In this case, the ‘true’ molecule is shown on the left, and 18 candidate molecules (from 100 runs) shown on the right. It is clear that all are close isomers, containing a methoxybenzoate moiety linked via a secondary amine to a pyrazole ring with a trifluoromethyl substituent.

As seen (Figure 2), while the model was run a hundred times, it predicted only 18 unique molecules, and each of them is a very close isomer. The closest one has a Tanimoto similarity of 0.95 to the true molecule when encoded with the TYPICAL fingerprint strategy^78^ as described next.

We ran a test set of 1350 separate molecules through the network using a single-blind strategy (mass peak lists of molecules not in the training or validation sets were provided by SS and were then tested by ADS). The concept of molecular similarity is quite elusive^84,85^, and the usual means of assessing it (an encoding of properties or molecular fingerprints followed by a similarity metric such as that of Jaccard or Tanimoto^86^) depends strongly on the encoding^78^, and is to some degree in the eye of the beholder. These comparisons are normally determined pairwise, though the parametrisation can be done using large cohorts of molecules^56^. For present purposes, we estimated similarity on the basis of what we refer to as the TYPICAL similarity^78^; this uses a set of encodings from rdkit (www.rdkit.org) (we used rdkit, atom pair, topological torsion, MACCS, Morgan and Pattern) followed by the Jaccard metric, and takes the values of whichever returns the largest (this was almost invariably the Pattern encoding).

Armed with the network’s estimates of the molecular structures and the then-disclosed identities, we used the above TYPICAL strategy^78^ to assess the Tanimoto similarity (TS) between the network’s best estimate and the true molecule. Of these, 1117 (82.7%) were precisely correct, 1338 had a TS exceeding 0.9 (99.1%) and only 2 (0.15%) had values of TS below 0.8. The data are given in Supplementary Table 1, and the structural closeness of some of the molecules with a TS of 0.9 is indicated in Figure 3. It is clear that a TS of greater than 0.9 does implies a molecule very close in structure to the correct molecule. Those shown give a clear indication of both the difficulty and the success of the task as they are indeed significantly complex structures. (Note that in this case 233 of the molecules contained S, of which 203 (91%) were nonetheless divined correctly.)

**Figure 3.**
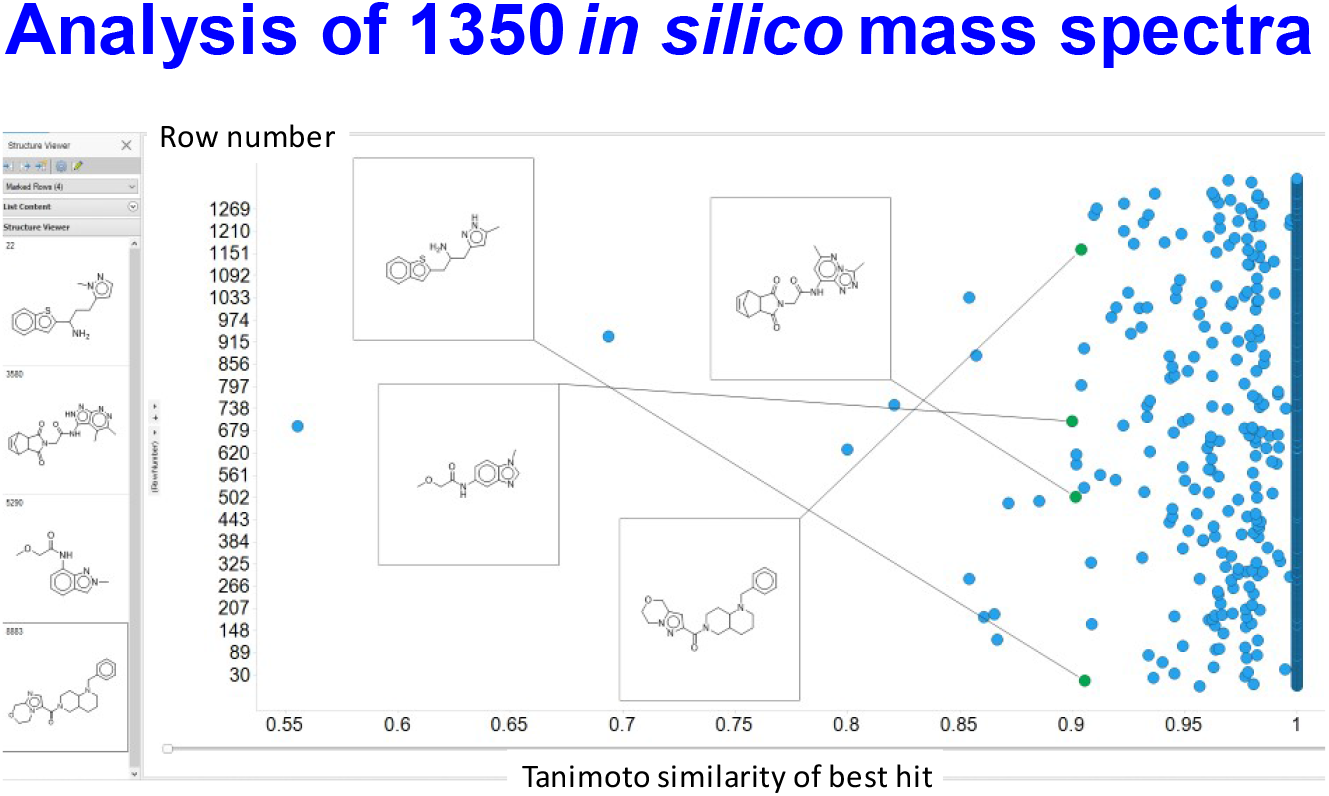
Analysis of a test set of predictions of the molecules behind 1350 *in silico*-fragmented mass spectral peaks. The peak lists were passed through the transformer after it had been trained and fine-tuned as described in methods. The analysis was done single-blind and the results fed back. The ‘closeness’ between the molecule estimated and the true molecule is given as the highest Tanimoto similarity based on six encodings. Four estimated molecules with a TS of ~0.9 are illustrated, together (at left) with the ‘true’ molecules. It is again obvious that they are extremely close structurally.

In a few cases, MassGenie failed to produce a molecule with the correct molecular formula. In this case, we were able to encode its best estimates in the latent space described in our VAE-Sim paper^56^, and move around the latent space until we found the closest molecule with the correct molecular formula. We ran 50 of these. Figure 4 illustrates the results for 10 molecules from those whose TS in Figure 3 was between 0.9 and 0.95, raising the average TS from 0.92 to 0.96, and producing structures that were clearly closer to the true one. The VAE-Sim strategy is thus a very useful adjunct to the direct Transformer approach.

**Figure 4.**
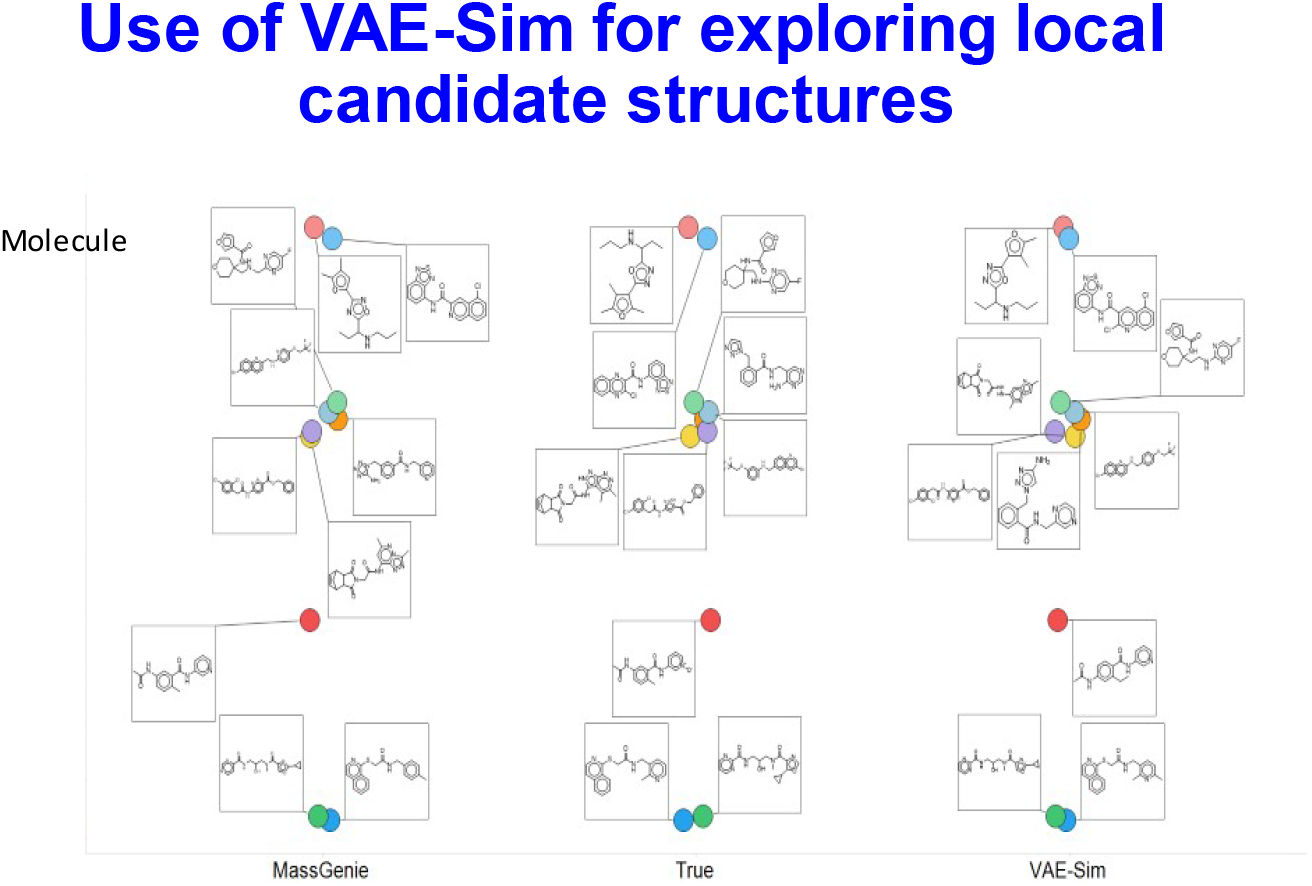
Local candidate structures generated by VAE-Sim. Based on 10 molecules taken from Fig 3, where the Tanimoto similarity between the best hit and the true molecule was in the range 0.9-0.95. VAE-Sim increases the number of candidates, in many cases improving their closeness to the true molecule.

Fig 5A further illustrates the use of VAE-Sim in cases where there is a mass difference between the known molecular ion and MassGenie’s predictions, both in terms of the rank order of cases where the mass difference is smallest (orange) or the Tanimoto similarity is greatest (blue). Fig 5B illustrates three examples in which VAE-Sim can effectively ‘recover’ the correct molecule by searching locally to the transformer’s top hits that just have the wrong mass.

**Figure 5.**
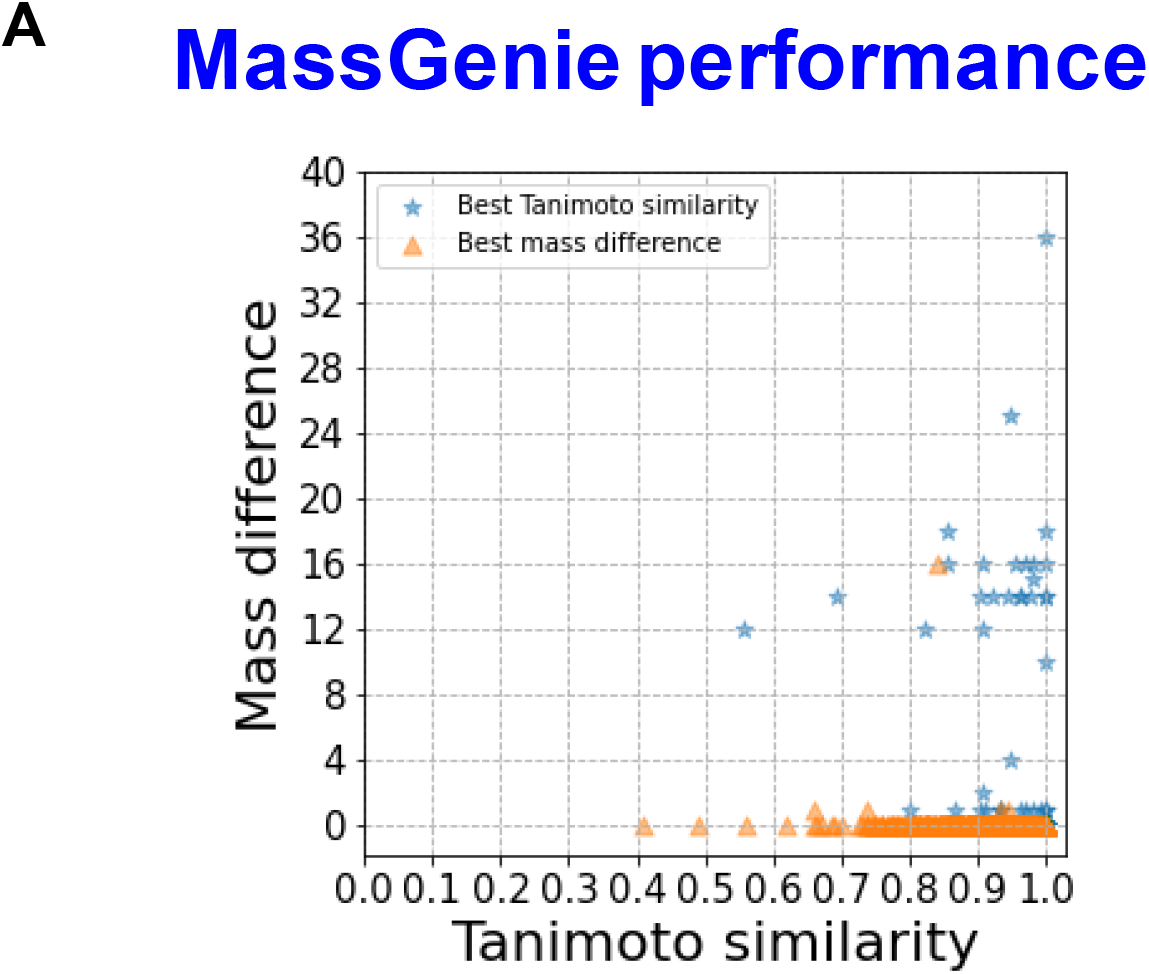
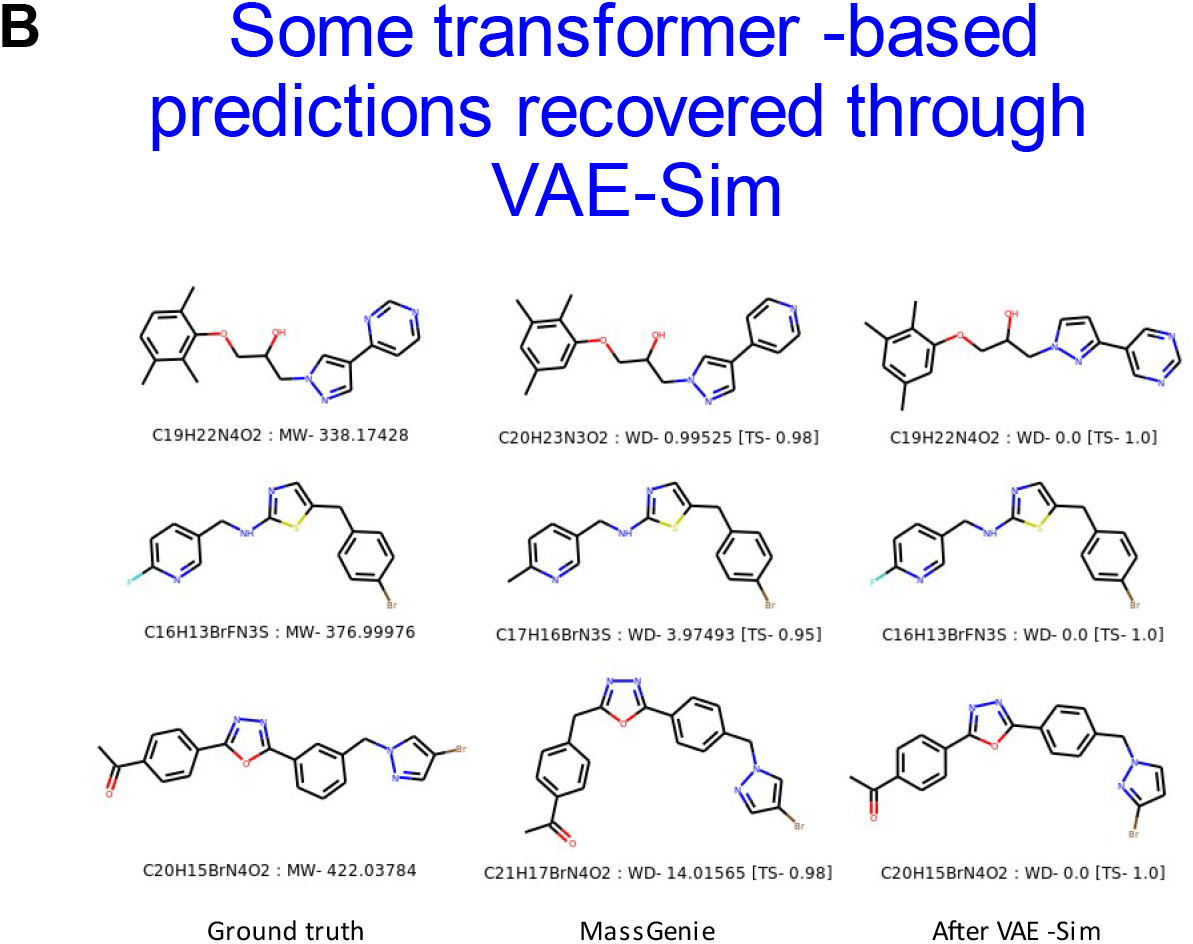
Local search where mass predictions are inaccurate. **A**. All examples from Fig 3 in which the transformer predicted molecules with slightly incorrect masses. VAE-Sim was used to search locally and generate further candidate structures that were round-tripped using FragGenie to produce mass spectra that could again be ranked in terms of TS to the known, true molecule given either the optimal mass difference or the optimal TS. **B**. Three examples showing the ground truth, the transformer’s best estimate, and the best (and accurate) prediction after the candidate pool of molecules was enhanced using VAE-Sim.

The number of fragments typically generated by FragGenie commonly exceeds those available in real electrospray mass spectra such as those available via MoNA^21,87^ or GNPS^21,87^ (and they are not identical in different databases). This said, generative deep learning methods of this type are sometimes capable of reconstructing whole entities (words or images) from partial, heavily masked inputs^88^, so we considered that it might be reasonable that a transformer trained solely on *in silico* fragments might nevertheless be able to make predictions from experimentally obtained mass spectra. We first used both purified standards of known structure and molecules assessed during experimental LC-MS(-MS) runs where spurious peaks can appear because of imperfect separations.

Our present LC-MS/MS method of the pooled human serum used^16^, based on that in^3^, shows 3009/3478 reproducible metabolites in ES^+^/ES^-^ mode, respectively, that appear in at least 50% of replicates and for which we have tandem mass spectra. Of these only 391/213 have confirmed entries in the mzcloud library (https://www.mzcloud.org/). Since the space of known and purchasable molecules far exceeds those for which we have experimental mass spectra, it is possible to assess those for which the proposed spectra are of purchasable compounds. It could thus be observed that experimental spectra of known compounds in serum often contained many spurious peaks not present in the standards. Consequently, we also used known standards that were run experimentally.

This experiment was again done single-blind, by which IR and MWM provided peak lists and knowledge of the molecular ions for molecules whose structures they had confirmed by running authentic standards or by other means. These are shown in Table 1, along with the performance of MassGenie. As with the interpretation of FragGenie peak lists, we obtained multiple candidates from each analysis. In practice, many of them possessed different SMILES representations but were actually of the same molecule. We therefore converted all the SMILES variants into a single canonical SMILES with its structure and assessed the most common. Figure 6 gives an example. As with FragGenie, the table thus uses as its top candidate the structures that were most frequently proposed. In some cases, MassGenie did not propose a structure with the correct mass ion.

**Table 1.**
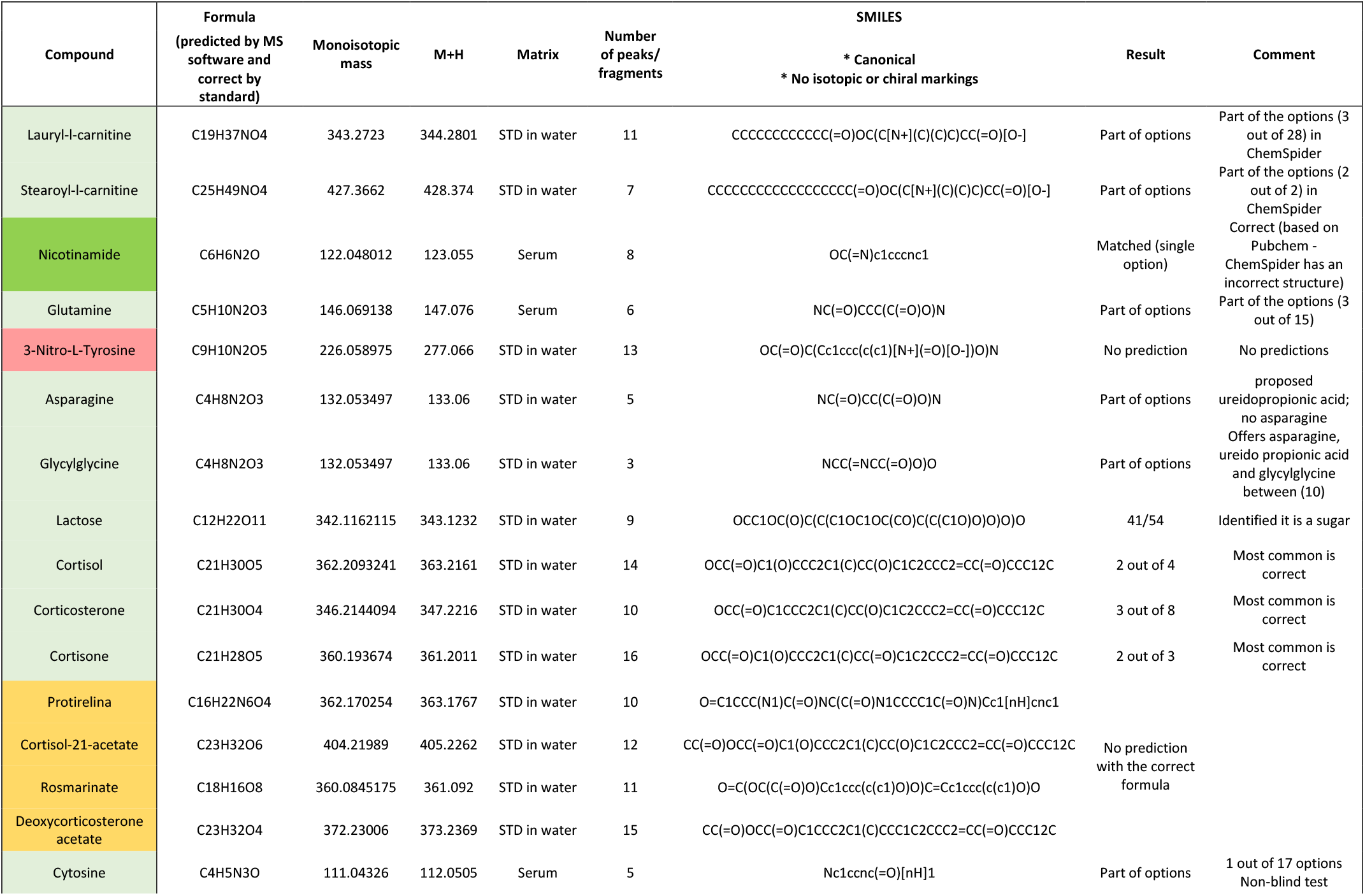
Predictions by MassGenie of molecular structures from experimental positive electrospray mass spectra, Molecules were provided single-blind to MassGeneie and the ranked outputs are recorded. 11 of the 16 molecules could be predicted.

**Figure 6.**
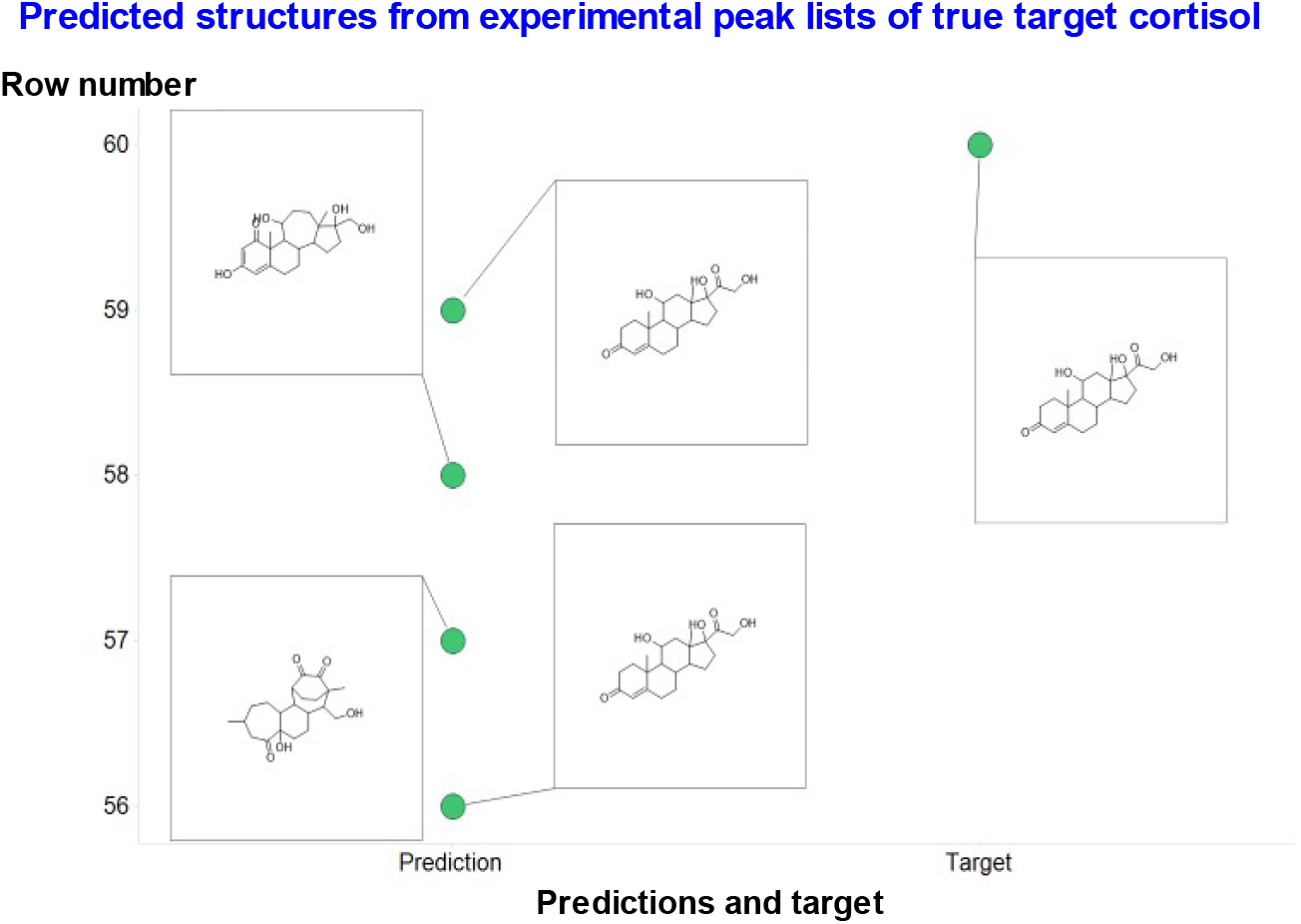
Illustration of the predictive power of MassGenie when presented with an experimental peak list (363.21631, 121.06477, 105.06985, 97.0648, 91.05423, 327.19522, 119.08553, 309.18469, 109.06478, 145.10121, 93.06989, 131.08549, 123.08039, 79.0542,143.08563) of the true molecule cortisol, along with its canonicalized best proposals. In this case 2/4 are correct (and to a chemist’s eye far more biologically plausible).

Overall, MassGenie performs very creditably, and where it does not one can begin to understand why. First, the experimental spectra often contain spurious peaks due to impurities, whereas those generated by FragGenie do not. Secondly, the number of experimental peaks is often really too low to allow a realistic chance of predicting a molecule much beyond those in the training set. Thirdly, there are obvious cases (e.g. long acyl chains) where FragGenie will produce a run of molecules differing by 14(.01565) mass units due to the serial loss of -CH2-groups, but the real electrospray mass spectra simply do not mirror this. This said, the fact that MassGenie, augmented where appropriate with VAE-Sim, can produce candidate molecules at all, including those that are not in any library, illustrates the massive power of generative methods of this type in attacking and potentially solving the enormous ‘mass-spectrum-to-structure’ problem of metabolomics and small molecule analytics.

## Discussion

A great power of modern methods of deep learning is that they do not need to be given explicit rules (though they or their human controllers may certainly benefit from them). The crucial point here, though, is that they are generative, i.e. in our case that they ‘automatically’ generate molecules in the form of SMILES strings; simple filters can allow the removal of invalid ones or those outside the appropriate mass range. VAE-Sim can be used to add to the list. Although future strategies will be able to incorporate the extensive existing knowledge of mass spectrometrists (e.g.^19,20,89^), the remarkable ability of our system to generate reasonably (and in a great many cases exactly) accurate molecules from candidate mass spectra, without any explicit knowledge of rules for the fragmentation of existing molecules and fragments, must be seen as striking. Note too that we are not directly using MS^n^ information based on parent ions beyond the protonated molecular ion at all. In contrast to methods designed to predict the presence of substructures alone, our system seeks to generate the entire molecule of interest, and can do so.

A 10×10 array of binary (black/white) pixels can take 2^100^ (~10^30^) forms^10^, but few of these represent recognisable or meaningful images. The existence of recognisable ‘features’ that may be extracted and combined to create meaningful images is what lies at the heart of the kinds of deep learning systems applied in modern, generative image processing^90^. In a similar vein, although the number of possible molecules is large, the number of meaningful (and chemically admissible) fragments or substructures from which they are built is far smaller^91^. Equivalently, an infinity of sentences may be constructed from a far smaller number of words, but only a subset of such sentences are meaningful and syntactically or semantically sound^92,93^. To this end, we do effectively cast the ‘mass-spectrum-to-structure’ problem as being equivalent to the kind of sequence-to-sequence language translation problem at which transformers excel, and we here apply them to this problem for the first time. Of the modern, generative methods of deep learning, transformers seem to be especially suited to these kinds of ‘language translation’ problems, and (for what we believe is the first time in this domain) this is what we have implemented here.

Modern methods in machine learning also recognise that it is possible to train deep networks to extract such features even from unlabelled image or textual inputs. In essence, our system relies similarly upon the use of generative methods with which we can embed molecules in a latent space from which they can be (re)generated. We can regress such latent spaces on any properties of interest. Commonly these latent spaces are used to predict the properties, but here we go in the other direction, where we use *in silico*-generated mass spectra to ‘predict’ the molecules from which they came. The key point is that this is now possible because we can generate them by the million (taking several hours per million on a V100- (not A100-) containing computer, depending on the depth of fragmentation requested). Implicitly, what these deep learning methods do is a form of statistical pattern recognition that effectively learns the rules of language and what letters are likely to be associated with what other letters and in what way. In our case, the ability to generate molecules from mass spectral peaks, i.e. fragments or substructures, shows that the transformer implicitly learns the rules of valency and which fragments may properly be ‘bolted together’ and how. This approach is in strong contrast (and in the opposite direction) to the generation of <u>experimental</u> mass spectra that (i) require access to known standards, (ii) are <u>highly</u> instrument-dependent, (iii) are time-consuming and expensive to acquire, and (iv) are consequently in comparatively short supply.

In the present work, we acquired a large number of molecules encoded as their SMILES strings, and used a gentler (and tuneable) modification of MetFrag^60,61^ that we call FragGenie to provide candidate peak lists and paired molecules whence they came. Although we anticipate the strategy to be generic, since this was a proof of concept we confined ourselves to ES+ spectra, and molecules with MW less than 500 and with SMILES encodings of fewer than 100 characters’ length. Its ability to predict unseen molecules from FragGenie lists was exceptional, with more than 99% being seen (after the single-blind code was broken) as having a Tanimoto similarity (using a suitable fingerprint encoding) of more than 0.9.

Of course, the real ‘proof of the pudding’ is how our predictions perform when experimental mass spectra are the inputs. In this case, some of the mass spectra contained noise peaks that the FragGenie spectra did not, and these meant that some molecules could not be learned; clearly the test data must bear a reasonable similarity to the training data if the system is to be able to generalise effectively.

Recently, Skinnider *et al*.^54^ applied a recurrent neural network-based generative model to attack a similar problem to that studied here, although they used a billion candidate structures and confined themselves to a test domain of some 2000 psychoactive substances. Although the approaches are not directly comparable, relatively few of their initial structures were accurate, although assessing them against the ‘structural priors’ in the billion molecules raised this to some 50%. Importantly, they recognised the value of constraining their candidate structures to those that generated the correct molecular ion, and they also showed the value of generative models for this problem.

Although we think that our strategy represents an excellent start, we should point out that this success did require what for an academic biology laboratory is a substantial computing resource – the DGX A100 8-GPU system we used allowed us to train a 400-million node network, which would in fact have represented a world record for the largest published deep learning network less than three years ago^72^. (This said, the <u>current</u> record, writing in June 2021, is more than three and almost four orders of magnitude greater^68,72,94^.) It is clear that the effectiveness of such networks will continue to increase, since how this potency scales with the size of the training dataset and of the network is broadly predictable^94–98^). We also recognise that MassGenie can be improved much further. First, although we have trained our system on millions of molecules, what is learned does tend to scale with the amount of material on which it is trained^99^. Indeed, there are already ~11 billion molecules in the ZINC database, and (as has already been done by Reymond and colleagues^41^) we can generate as many more (in both type and number) as we like *in silico;* using these in further training, probably with larger encoding networks, will require more computer power, but this is evidently going to become available (as noted above, e.g. in the training of GPT-3^68^ and others^100^). Secondly, the rules we have used in the *in silico* fragmentation are extremely crude and rudimentary; they can certainly bear refining or replacing with some of the more knowledge-based *in silico* fragmentation methods available in commercial software (which would have to have an API to admit the generation of large numbers of SMILES-spectrum pairs, as here). Thirdly, we have not used any knowledge of isotope patterns and the like, relying solely on exact masses to discriminate the likely atoms in a fragment. Fourthly, we have used SMILES strings as our molecular encoding; a move to graphical representations^101–106^ is likely to prove beneficial. Fifthly, we have yet to apply equivalent strategies to negative ionisation spectra and to larger molecules, and to a far greater number of molecules containing P and S. Sixthly, by using canonical SMILES we ignore – and thus cannot discriminate – any stereoisomerism. Finally, we did not really explore at all the many flavours of deep network being developed to learn, store and transform information^107,108^. Although this is a very fast-moving field, those generative methods based on transformers^70,109–111^, as we use in the spectrum-to-structure problem for the first time here, do presently seem particularly promising. In line with the title of the original transformer paper^70^, we might conclude by commenting that “accurate fragmentation is all you need”.

## Supporting information

Test data from FragGenie plus predictions as described in text

## Conflict of interest statement

The authors declare that they have no conflicts of interest.

## Author contributions

ADS wrote the transformer code, NS wrote FragGenie, SS wrote and applied VAE-Sim. NS and SS constructed the datasets. IR and MWM ran experimental mass spectra and provided peak lists and cheminformatic analyses. DBK conceived the project, obtained funding for it, and wrote the first draft of the manuscript. All authors contributed to further drafting and editing, and approved the final version.

## Acknowledgments

ADS was funded via the University of Liverpool. SS and DBK are funded as part of the EPSRC project SuSCoRD (EP/S004963/1), partly sponsored by Akzo-Nobel. DBK is also funded by the Novo Nordisk Foundation (grant NNF20CC0035580). The DGX-A100 was purchased as part of a BBSRC infrastructure grant to the University of Liverpool GeneMill. We thank Pete Balshaw (Scan Computers) and Cliff Addison (University of Liverpool) for their help in its specification and installation.

## Code availability

The code will be made available via links in the Supplementary Information upon publication in a peer-reviewed journal. Note, however, that the trained transformer model occupies 1.2Gb.

## Supplementary Information

File test_set_data_analysis.xlsx representing the peak lists and molecules shown in Figure 3.

## Notes

### Competing Interest Statement

The authors have declared no competing interest.

